# Transcriptional heterogeneity in cancer-associated regulatory T cells is predictive of survival

**DOI:** 10.1101/478628

**Authors:** Nicholas Borcherding, Kawther K. Ahmed, Andrew P. Voigt, Ajaykumar Vishwakarma, Ryan Kolb, Paige Kluz, Gaurav Pandey, Katherine N. Gibson-Corley, Julia Klesney-Tait, Yuwen Zhu, Jinglu Lu, Jinsong Lu, Xian Huang, Jinke Cheng, Song Guo Zheng, Xuefeng Wu, Yousef Zakharia, Weizhou Zhang

## Abstract

Regulatory T cells (Tregs) are a population of T cells that exert a suppressive effect on a variety of immune cells and non-immune cells. The suppressive effects of Tregs are detrimental to anti-tumor immunity. Recent investigations into cancer-associated Tregs have identified common expression patterns for tumor-infiltration, however the functional heterogeneity in tumor-infiltrating (TI) Treg is largely unknown. We performed single-cell sequencing on immune cells derived from renal clear cell carcinoma (ccRCC) patients, isolating 160 peripheral-blood (PB) Tregs and 574 TI Tregs. We identified distinct transcriptional TI Treg cell fates, with a suppressive subset expressing CD177. We demonstrate CD177^+^ TI-Tregs have preferential suppressive effects in vivo and ex vivo. Gene signatures derived the CD177^+^ Treg subset had superior ability to predict survival in ccRCC and seven other cancer types. Further investigation into the development and regulation of TI-Treg heterogeneity will be vital to the application of tumor immunotherapies that possess minimal side effects.

---

Regulatory T cells (Tregs) are a population of T cells that exert a suppressive effect on a variety of immune cells and non-immune cells^1^. Similar to the role in peripheral tolerance, the suppressive effects of Tregs are detrimental to anti-tumor immunity^1^. The prognostic value of *FOXP3*, the master regulator of Tregs, is highly variable^2–6^. Recent investigations have identified common expression patterns for tumor-infiltrating Treg (TI-Treg)^7–9^; however, the heterogeneity of TI-Tregs at the functional levels is largely unknown across different cancer types. We performed single-cell RNA sequencing (SCRS) on immune cells derived from renal clear cell carcinoma (ccRCC) patients, isolating 160 peripheral-blood (PB) Tregs and 574 TI-Tregs. We also analyzed a published SCRS datasets containing 634 TI-Tregs from human hepatocellular carcinomas (HCC). We identified distinct transcriptional cell fates in both ccRCC and HCC, with the suppressive subsets of TI-Tregs expressing a common gene signature including *CD177* as the most upregulated gene. We further demonstrate CD177^+^ Tregs have preferential suppressive effects across human and mouse. Gene signatures derived from CD177^+^ Tregs had superior predictability for patient survival in several cancer types, outperforming other reported Treg signatures. Further investigation into the development and regulation of TI-Treg heterogeneity will be vital to the application of tumor immunotherapies with potential minimal side effects.

In order to investigate the heterogeneity and dynamics of TI-Tregs, we performed SCRS on peripheral and TI immune cells from three ccRCC patients. ccRCC tumors are responsive to immune checkpoint blockade despite low mutational loads, indicative of microenvironmental involvement^10^. A total of 25,672 immune cells were sequenced, with 160 PB and 574 TI-Tregs isolated using the expression of *FOXP3* and CD25 (*IL2RA*) (Figure 1a-b). To identify differentially-expressed genes and markers of TI-Tregs, we compared TI-*versus* PB-Tregs expression (Figure 1c). A complete list is available in Supplemental Table 1. The only two genes with no expression in any PB Tregs were *NR4A1* (51.4%) and *CD177* (20.2%). A summary is shown for the top eight upregulated (Figure 1d) or downregulated (Figure 1e) in TI-Tregs shown as log2-fold change (LFC).

**Figure 1.**
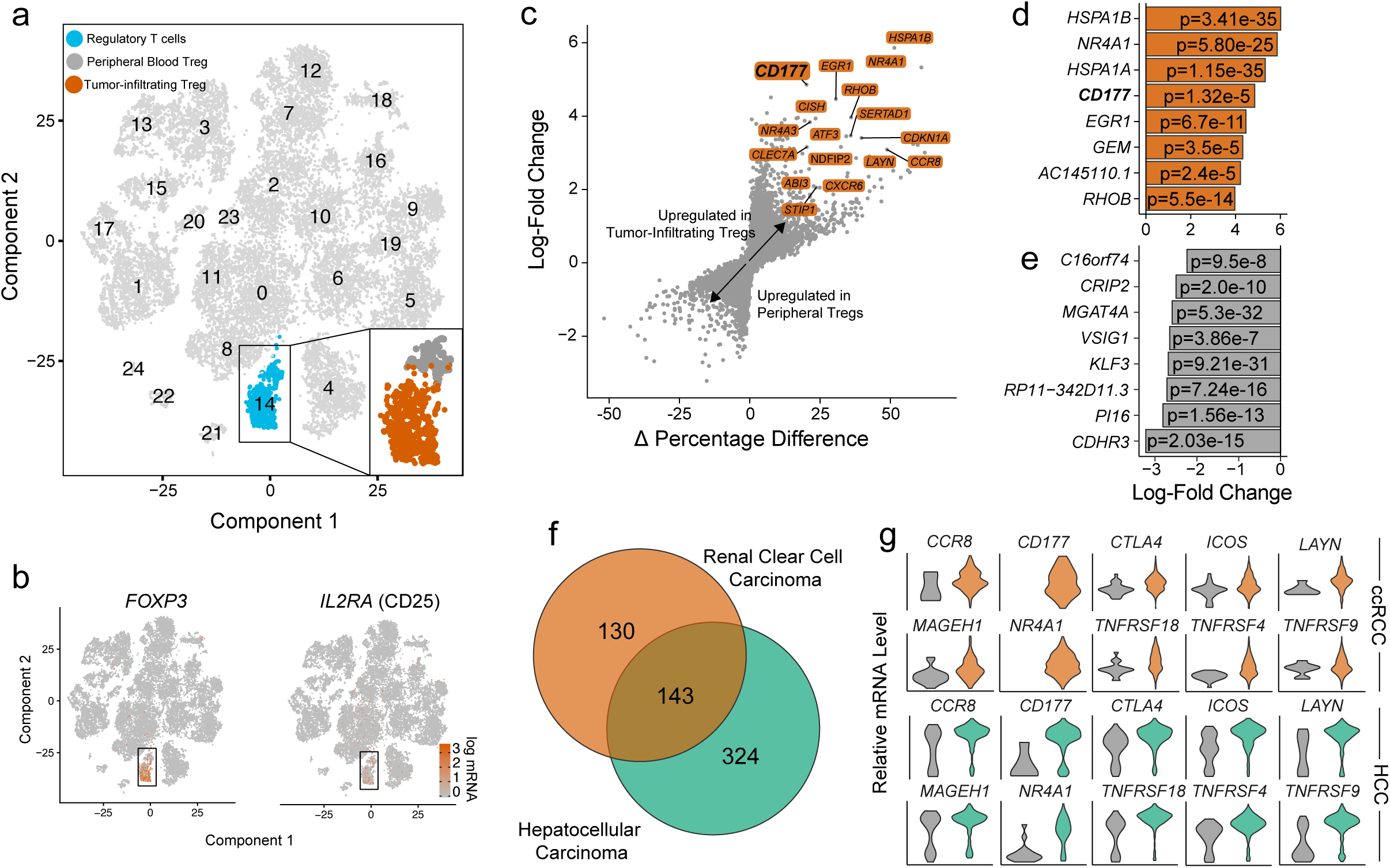
TI-Tregs display distinct expression program compared to PB controls in ccRCC SCRS dataset. (a). tSNE projection of immune cells from three ccRCC patients with normal PB cells (n=13433) and TI cells (n=12,239). Treg population (blue) was isolated and separated as TI (orange) *versus* PB Tregs (grey). (b). tSNE projection with highlighted expression of Treg markers, *FOXP3* and *IL2RA* (CD25). (c). Differential gene expression analysis using the log2-fold change expression versus the difference in the percent of cell expressing the gene comparing TI versus PB Tregs (Δ Percentage Difference). Genes labeled have log2-fold change > 1, Δ Percentage Difference > 20% and adjusted P-value from Wilcoxon rank sum test < 0.05. (d). Top eight upregulated genes by log2-fold change in TI-Tregs with adjusted P-value < 0.05. (e). Top eight downregulated genes by log2-fold change in TI-Tregs with adjusted P-value < 0.05. (f). Comparison of differential genes in TI-Tregs in ccRCC (orange) and HCC (green) compared to PB Tregs. Significant genes were defined as log2-fold change > 1 or < −1 with adjusted P-values < 0.05. (g). Relative mRNA level of Treg markers in PB (grey) and TI-Tregs in ccRCC (top) and HCC (bottom).

We also analyzed a recently-published single-cell profile of T cells in HCC ^9^, comprised of flow-sorted T cells including Tregs from PB, normal liver parenchyma, a transitional zone near the tumor, and TI-Tregs. After clustering (Supplemental Figure 1a-b), we performed the same differential gene analysis comparing TI-Tregs *versus* PB-Tregs (Supplemental Figure 1c). We found a total of 273 differential genes in our ccRCC-infiltrating Tregs, 467 differential genes in the HCC-infiltrating Tregs, and 143 shared differential genes (Figure 1f, Supplemental Figure 1c). Of note, we corroborated previous reports of increased *CCR8, LAYN*, and *MAGEH1* expression in TI-Tregs (Figure 1f-g)^7–9^. The gene with the highest expression in ccRCC and HCC TI-Tregs was *CD177*. A similar pattern of upregulated genes, including *CD177*, was seen in pooled Tregs in breast, colorectal, and lung cancers (Supplemental Figure 1d-e).

Recent studies started to imply functional heterogeneity of FOXP3^+^ Tregs in PB^11^ as well as colorectal cancer^12^ and glioma^13^, based on genomic studies with pooled Tregs. Using SCRS, we were able to investigate the dynamic transcriptomic processes of individual Tregs. Using the Monocle 2 algorithm^14^, we constructed a manifold using all the Tregs in ccRCC (Figure 2a) and HCC (Supplemental Figure 2a). This technique orders the cells by expression pattern to represent distinct cellular fates, with the ordinal construction creating a pseudo-time variable that allows us to investigate changes in gene expression during the infiltration process. We observed a bifurcated architecture of TI-Tregs in ccRCC (Figure 2a) and HCC (Supplemental Figure 2a), implying two distinct cell fates. We also found a number of genes with significant roles in the manifold ordering (Figure 2b), with complete results available in Supplemental Table 2. A similar profile of genes was involved in manifold construction of the HCC Tregs with reduced separation between TI *versus* PB Tregs in several genes including *CTLA4* and *CCR8* (Supplemental Figure 2b). In contrast, other genes like *CCL20* and *CD177* maintained distinct separation in expression from TI *versus* PB Tregs (Supplemental Figure 2b).

**Figure 2.**
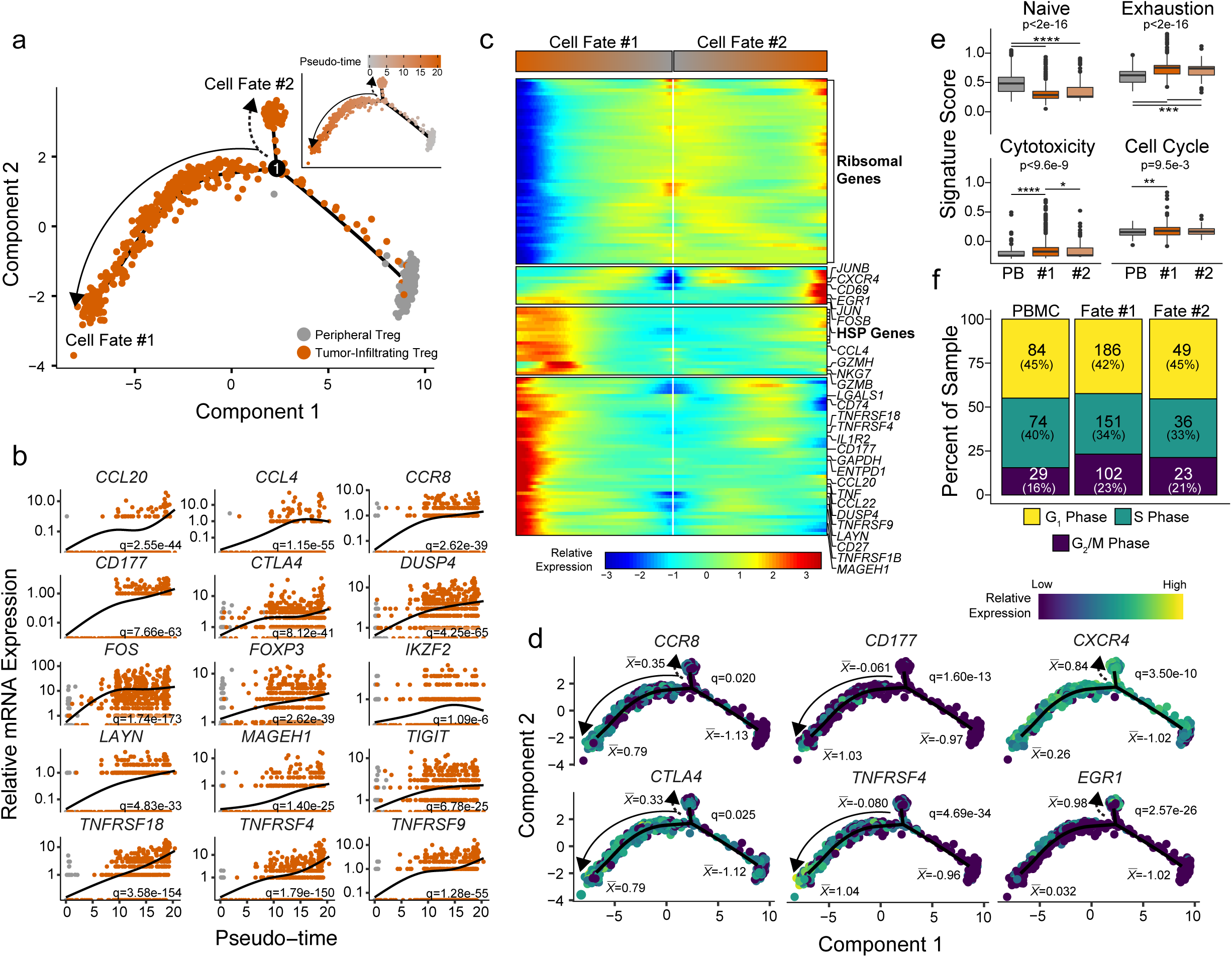
Bifurcation in the transcriptional state of TI-Tregs reveal a more suppressive cell fate. (a). Trajectory manifold of Tregs from the renal clear cell carcinoma using the Monocle 2 algorithm, solid and dotted line represent distinct cell trajectories/fates defined by SCRS expression profiles. (b). Pseudo-time projections of transcriptional changes in immune genes based on the manifold. Significance based on differential testing by site of origin which was also used to generate pseudo-time and adjusted for multiple comparisons. (c). Expression heatmap of significant (Q < 1e-6) genes based on branch expression analysis comparing the two TI cell fates and were used in the ordering of the pseudo-time variable. (d). Cell trajectory projections of transcriptional changes in immune genes based on the manifold. Significance based on differential testing between the first and second cell fates of TI-Tregs. 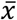 denotes the scaled mean mRNA levels of each pole of the manifold. (e). SCRS gene signature analysis of the poles of the trajectory manifold. P-value based on one-way ANOVA with individual comparisons corrected for multiple hypothesis testing using the Tukey HSD method. * P < 0.05, ** P <0.01, *** P <0.001, **** P < 0.0001. (f). Results of the cell cycle regression analysis of single cells for each cell fate using the Seurat R package.

Next, to understand the branching structure of the manifold, we performed differential gene expression analysis. Markers of immune regulation had increased expression in TI-Tregs of the first cell fate (CF1) compared to the Tregs of the second cell fate (CF2, Figure 2c, lower two clusters). In contrast, the CF2 had a maintenance of ribosomal gene expression (Figure 2c). We observed three trends in gene expression between the TI-Tregs cell fates: non-specifically increased, like *CCR8* and *CTLA4*; increased in CF1, like *CD177* and *TNFRSF4*; and increased in CF2, like *CXCR4* and *EGR1* (Figure 2d). To investigate the potential functional difference between the cell fates, we performed gene signature analysis (Figure 2e). TI-Tregs had a reduction in the naïve T cell signature and an increase in T cell exhaustion signature compared to PB Tregs in both ccRCC (Figure 2e, upper row) and HCC (Supplemental Figure 2d). In contrast, the cytotoxicity and cell cycle signatures were significantly higher in CF1 Tregs (Figure 2e, lower row). We assigned cell phases to single cells, finding an increased percentage of G_2_M phase Tregs in in CF1 (Figure 2f). This increase in G_2_M was not demonstrated in the CF1 Tregs in the HCC dataset, indicative of cancer-specific properties of TI-Tregs (Supplemental Figure 2d-e). We also observed a clonal expansion in TI-Tregs across all ccRCC patients (Supplemental Figure 3a-c), in line with previous observations^8,9,15–17^. In addition, we noticed that the clonal expansion of the T cell receptor repertoire was enriched in the Tregs of CF1 (Supplemental Figure 3d).

**Figure 3.**
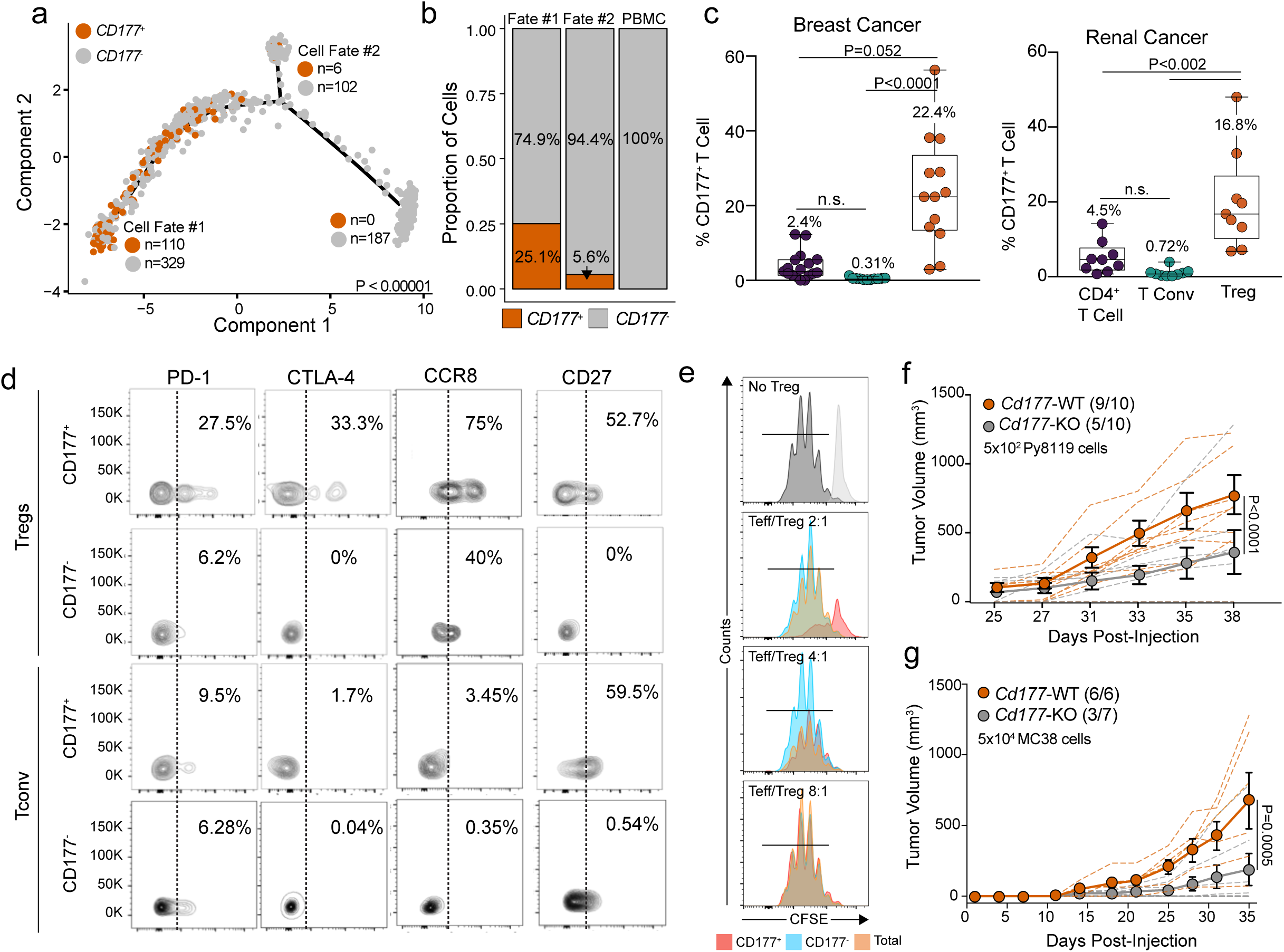
CD177 is a marker a suppressive subpopulation of TI-Tregs. (a). Trajectory manifold of Tregs from the ccRCC Tregs with the number of *CD177*^+^ and *CD177*^-^ Tregs for each respective cell fate. Significance based on X^2^ testing comparing the three poles of the manifold. (b). Proportional distribution of *CD177*^+^ Tregs by cell fate across the manifold. (c). Percentages of CD177^+^ cells relative to different CD4^+^ TILs in breast cancer (n=13-18) and renal cancer (n=8) showing Data are presented as mean ± SD with median values presented in the figure. (d). Representative flow cytometry data for select marker expression by CD177^+^ or CD177^-^ Tregs (CD3^+^CD4^+^CD25^+^CD127^low^/FoxP3^+^) or Tconv (D3^+^CD4^+^CD25^-^/FoxP3^-^) isolated from breast cancers. n=2-3. (e). CD177^+^ TI-Tregs are suppressive and inhibit CD8 T cell proliferation. Total, CD177^+^, or CD177^-^ TI-Tregs were purified from fresh human breast cancer specimens (combined from 3 patients) using flow cytometry and co-cultured with naïve CD8 T cells from PBMC for *ex vivo* suppression assay at the indicated ratios. (f). Py8199 tumor growth is significantly reduced in *Cd177-*KO mice compared to WT, P < 0.0001 (two-way ANOVA) in mice challenged with 5×10^2^ cells per inoculation, n=10 bilateral tumors. Numbers in parenthesis equates to the number of mice which developed palpable tumors/total mice inoculated. Data are presented as mean ± SEM. (g). MC38 tumor growth is significantly reduced in *Cd177-*KO mice compared to WT, P = 0.0005 (two-way ANOVA) in mice challenged with 5×10^4^ cells per inoculation. Numbers in parenthesis equates to the number of mice which developed palpable tumors/total mice inoculated. Data are presented as mean ± SEM.

The increased expression of *CD177* in both ccRCC and HCC Tregs compared to PB Tregs (Figure 1g) and the specificity towards the suppressive CF1 (Figure 3a-b) led us to further determine the functional impact of CD177^+^ Tregs. CD177 is a glycosylphosphatidylinositol-linked surface protein that is expressed on neutrophils^18^ and has been used as a biomarker for myeloproliferative diseases^19^. Our interest in CD177 is long standing and we have previously shown that CD177 plays a role in neutrophil viability^20^. We performed flow cytometry on tumor-infiltrating T lymphocytes (TITL) and demonstrated that a majority of CD177^+^ cells were Tregs (Supplementary Figure 4a-b). There were 22.4% CD177^+^ Tregs among TI-Tregs in breast cancer and 16.8% in renal cancer (Figure 3c). There were negligible percentages of conventional CD4^+^ T cells (Tconv) expressing CD177 in breast and renal cancers (Figure 3c). Additionally, we developed an immunohistochemical (IHC) protocol (Supplemental Figure 4c-d), finding CD177^+^ lymphocytic infiltration was enriched in the colon, lung, liver, and prostate. In addition to the single-color IHC, we further characterized the lymphocytic infiltration in breast tumors using dual-IHC staining, identifying CD177 and FOXP3 double-positive cells (Supplemental Figure 4e). Further analysis revealed that CD177^+^ Tregs had larger pools of cells expressing PD-1, CTLA-4 and CCR8 when compared to CD177^-^ Tregs, suggesting an active and suppressive phenotype (Figure 3d). CD27, an activation marker, exhibited a similar pattern for CD177^+^ Treg or Tconv cells (52.7% vs 59.5%), but was lost in CD177^-^ Tregs (Figure 3d). We were also able to isolate a fraction of CD4^+^ conventional T cells (Tconv) with CD177 expression failed to show a clear differential pattern compared to CD177^-^ Tconv cells (Figure 3d), indicative of a Treg-specific phenotype related to CD177 expression.

**Figure 4.**
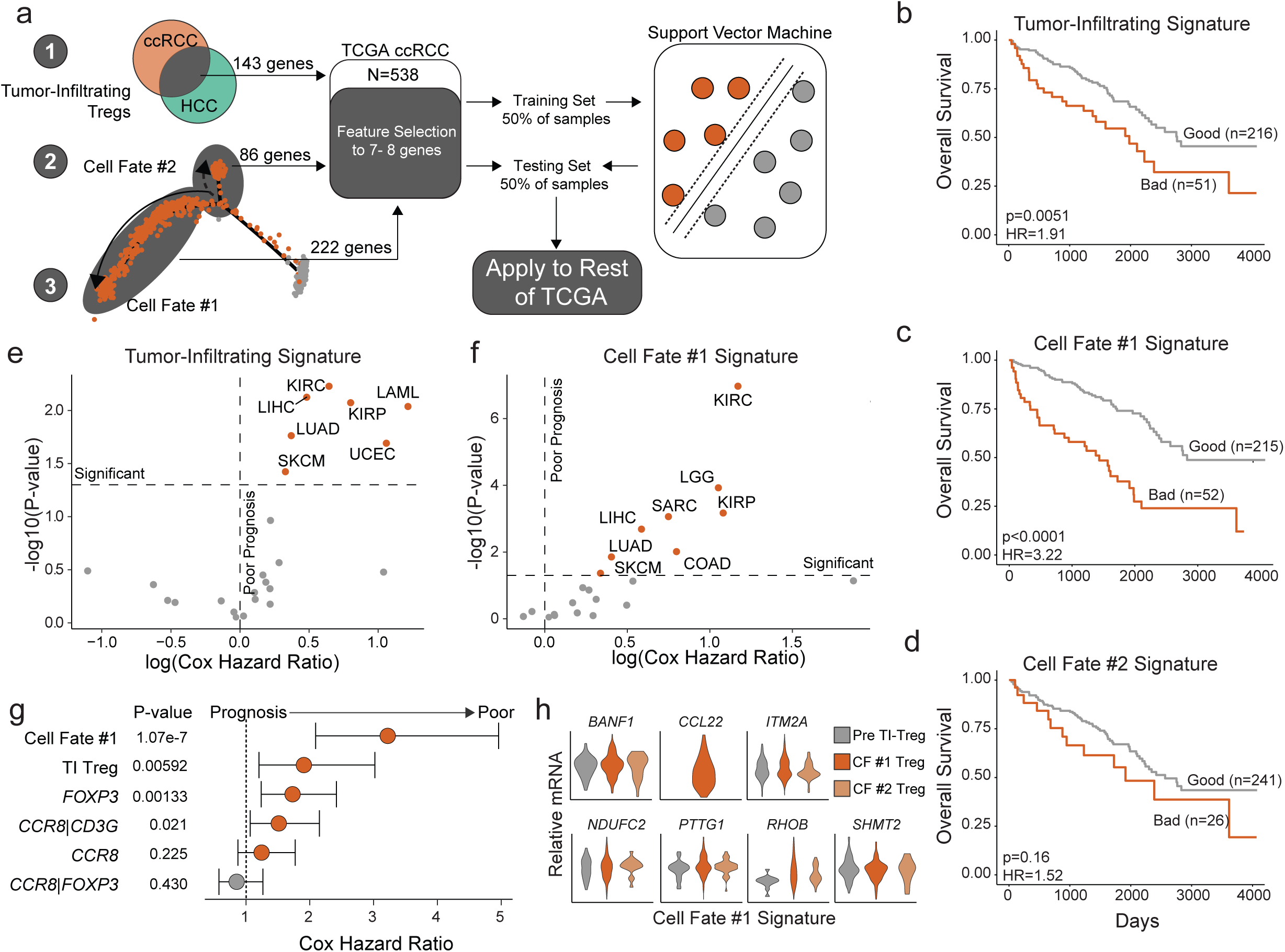
Improved prognostic prediction associated with signature from suppressive Treg cell fate. (a). Schematic of signature development using feature selection from: 1) 143 differential genes of TI-Treg in ccRCC and HCC, 2) 86 genes differentially expressed in CF2, and 3) 222 genes differentially expressed in CF1 using the ccRCC (n=538) dataset from the TCGA. (b-d). Kaplan-Meier curves for overall survival in ccRCC using the TITL gene signature (b), in ccRCC using the CF1 Treg gene signature (c), and the CF2 Treg signature (d). P-value based on log-rank test and hazard ratio (HR) based on Cox proportional hazard regression. (e-f). Overall survival prediction with Cox proportional hazard ratio and −log10(P-value) based on log-rank testing across the 24 largest TCGA datasets using the TI-Treg signature (e) and the CF1 signature (f). (g). Prognostic prediction for SCRS Treg signatures compared to other proposed signatures for TI-Tregs. Hazard ratios, 95% confidence intervals, and P-values derived from Cox proportional hazard regression modeling. (h). Relative mRNA violin plots of the CF1 signature based on the transcriptional trajectory state across the 160 PB-Tregs and 574 TI-Tregs.

In order to confirm the immunosuppressive property of CD177^+^ Tregs, we performed *in vitro* proliferation and suppression assays from patient-derived TITL cells (Figure 3e, Supplemental Figure 5a). Results showed that CD177^+^ Tregs had a 4-fold greater suppressive effect on CD4^+^ effector T cells when plated at a 2:1 ratio compared to CD177^-^ Tregs (73% versus 16%) (Figure 3E, Supplemental Figure 5A). The scarcity of CD177^+^ tumor infiltrating Tregs made it challenging to repeat these experiments. However, given that the experiments were carried out using Tregs pooled from three individual breast cancer tissues, results can be considered biologically representative.

We used *Cd177*-KO mice and their WT littermate controls to verify the suppressive phenotype of CD177^+^ Tregs. After titrating injections to identify optimal cell numbers to facilitate immune-based tumor control of Py8119 cells, a breast cancer model, we found an optimal dose of 500 cells. We observed a clear phenotype in regard to tumor growth in *Cd177-*KO mice *versus* WT, where KO mice rejected tumor inoculation (5 of 10 total) and showed significantly slower tumor growth (Figure 3f). The same increase in tumor rejection (4 of 7) and slower tumor growth was seen in the syngeneic graft of MC38 colorectal cells in *Cd177*-KO *versus* WT controls (Figure 3g). To confirm that the observed phenotype is, in part, due to the suppressive action of CD177^+^ Tregs in WT mice, we performed an *in vitro* immunosuppression assay using Tregs sorted from splenocytes of tumor-bearing *Cd177-*KO and WT mice. Tregs isolated from WT mice were more suppressive of CD4^+^ and CD8^+^ T cell proliferation compared with *Cd177*-KO Tregs (Supplemental Figure 5b).

As our data suggests the transcriptional and functional difference in a suppressive subset of tumor-infiltrating Tregs across several cancers, we hypothesized that gene signature development from SCRS data would provide improved prognostic ability. We performed feature selection to identify sets of genes most associated with overall survival for: 143 differentially-expressed genes of TI-Treg, 222 genes differentially-expressed in CF1, and 86 differentially-expressed genes in CF2 using the ccRCC dataset from the Cancer Genome Atlas (TCGA, Figure 4a). Using 50% of the ccRCC samples as a training set, we trained supervised support vector machines to discriminate between survival outcomes. Applying these signatures to the remaining 50% of samples, we showed the TI (Figure 4b) and CF1 signatures (Figure 4c) discriminated prognosis in ccRCC, but not the CF2-based signature (Figure 4d). Despite both the TI and CF1 signature significantly predicting poor overall survival in roughly 20% ccRCC patients, CF1 signature has a superior ability to discriminate prognosis (hazard ratio of 3.22 *versus* 1.91) (Figure 4b-c).

The superior discrimination of CF1 Treg signature compared to the TI-Treg signature was observed across a number of cancers. We applied the ccRCC-signatures across the 24 largest TCGA datasets, finding that the TI-Treg signature significantly separated prognostic groups in 7 cancer datasets, with hazard ratio range limited to 1.39 to 3.37 (Figure 4e). In contrast, the CF1 Treg signature discriminated prognostic groups in 8 cancer types, with a larger hazard ratio range of 1.4 to 5.5 (Figure 4f). The greater number and range of significant predictions by cancer was also seen when comparing disease-specific survival in CF1 Treg *versus* TI-Treg signature (Supplemental Figure 6a-b). CF2 Treg signature failed to discriminate groups based on overall survival in any TCGA dataset (Supplemental Figure 6c). Both the CF1 Treg and the TI-Treg signature separated prognostic groups in immune-checkpoint-inhibitor responsive melanoma (SKCM) and lung adenocarcinoma (LUAD) (Figure 4e-f). The CF1 Treg signature separated prognostic groups in sarcoma (SARC), low-grade glioma (LGG) and colon adenocarcinoma (COAD). Both the CF1 Treg and TI-Treg signature had larger hazard ratios than previously identified tumor Treg markers, like *FOXP3*, and ratio-based signatures *CCR8*:*FOXP3* or *CCR8*:*CD3G* (Figure 4g)^7,8,21^. It should be noted that feature selection of the CF1 did not include CD177 into the gene signature, likely due to the expression of CD177 in epithelial tissue (Figure 4h, Supplementary Figure 4).

In conclusion, although infiltration of FoxP3^+^ Tregs are thought to suppress antitumor immune response broadly, our work demonstrates transcriptional and functional heterogeneity with TI-Tregs. Using SCRS of 734 Tregs, we showed two distinct transcriptional fates of TI-Tregs (Figure 2a). Although a transcriptional analysis of pooled TI-Treg was previously found incidentally,^8^ we herein identified CD177 as a marker of a suppressive subset of Tregs with superior ability to inhibit effector T cell populations and tumor growth in both breast and colorectal models. Comparing prognostic ability for Treg gene signatures, we demonstrated the superior outcome prediction for the suppressive CF1 signature in demarcating prognostic groups in ccRCC and across the TCGA compared to the TI-Treg gene signature or CF2 Treg signature (Figure 4). Our study displays the potential of using SCRS to focus investigation of specific immune subsets and potentially identify new therapies or prognostic markers. Further understanding of the development and regulation of this suppressive subset of TI-Tregs may be crucial in the development of cancer immunotherapies with minimal autoimmune side effects.

## Authors’ Contributions

Conception and design: W.Z., N.B. K.K.A, X.W., Y.Z.

Development of methodology: N.B., K.K.A., P.K., K.G.C., A.V.

Acquisition of data: K.K.A., N.B., R.K., A.V., P.K., G.P., K.G.C., J.K.T., Y.W.Z., J.L., X.H., X.W., W.Z.

Analysis and interpretation of data: N.B., K.K.A., R.K., A.V., X.W., Y.W.Z., W.Z.

Writing, review, and/or revision of the manuscript: N.B., K.K.A, R.K., Y.W.Z, S.G.Z, Y.Z., W.Z.

Study supervision: X.W., Y.Z., W.Z.

## Acknowledgments

We thank Breast Molecular Epidemiologic Resource (BMER led by Dr. Sonia Sugg) and Tissue Procurement Core at the University of Iowa/Carver College of Medicine for providing breast cancer tissues; TCGA for providing breast cancer, melanoma and other cancer RNA-seq datasets and clinical parameters; Comparative Pathology Laboratory from Department of Pathology University of Iowa/Carver College of Medicine for developing CD177 IHC protocol.

