## Supplementary material for "Transcriptional heterogeneity in cancer-associated regulatory T cells is predictive of survival"

### Materials and Methods

*Patient Recruitment*

The current study was approved by the University of Iowa Institutional Review Board (IRB) and conducted under the Declaration of Helsinki Principles. De-identified renal patients were recruited by Dr. Yousef Zakharia in the Department of Internal Medicine at the University of Iowa. These tissues were also collected from patients undergoing surgical resection after informed consent and were supplied as de-identified samples to Dr. Zhu laboratory with approved IRB approval. Additional human specimens were collected from the Department of Breast Surgery, Renji Hospital, Shanghai Jiao Tong University School of Medicine in a de-identified manner to the Dr. Wu laboratory for cell preparation and T cell suppression assay, approved by the University Human Subject Protection Committee.

*Single-cell RNA sequencing*

Viable (Hoechst$^{-}$), immune (CD45$^{+}$) single-cell suspensions generated from three ccRCC tumor samples and peripheral blood were sorted on a FACS ARIA sorter (BD Biosciences) for lymphoid and myeloid lineages. The cells were sorted into ice-cold 1x PBS + 0.04% non-acetylated bovine-serum albumin (New England BioLabs, Ipswitch, MA). Sorted cells were counted and viability as assessed using the MoxiGoII counter (Orflo Technologies, Ketchum, ID). Single cells were re-suspended at 1000 cells/µl with cell viability greater than 90%. Sequencing was performed using the Chromium (10x Genomics, Pleasanton, CA) and Illumina (San Diego, CA) sequencing platforms. Amplified cDNA was used to construct both 5’ expression and VDJ sequencing libraries.

Pooled libraries were run on sperate lanes of a 150 based-paired, paired-end, flow cell using the Illumina HiSeq 4000 in the University of Iowa Genomics Division. Basecalls were converted into FASTQs using the Illumina bcl2fastq software by the University of Iowa Genomics Division. FASTQ files were aligned to human genome (GRCh38) using the CellRanger v2.2 pipeline as described by manufacturer[1]. Individual cells were quality checked for total expression and percentage of mitochondrial reads. In order to reduce reads from potential multiple cells, cells with greater than 5,000 unique genes were filtered out of the data. After processing, clustering was performed using the Seurat R package (v2.3.4), correcting for patient variability using the canonical correlation process[2,3]. Dimensional reduction to form the tSNE plot utilized the top 20 calculated dimensions and a resolution, or granularity of the clusters, of 1.2. Differential gene expression analysis was performed using the Wilcoxon rank sum test for significance comparing tumor-infiltrating *versus* peripheral blood Tregs. Cell trajectory manifolds and pseudo-time estimates utilized the Monocle R package (v2.8.0) and the reverse graph embedding machine learning algorithm[4]. Single-sample gene set enrichment analysis utilized singleR R package (v0.2.0)[5] and cell cycle regression for individual Tregs was performed in the Seurat R Package, as previously described[6]. The same processing and quality control procedures were applied to count-level data from GSE98638[7].

Bulk RNA sequencing Treg Data

Raw expression data for GSE89225[8] and PRJEB11844[9] were downloaded from the NCBI Sequence Read Archive and the European Nucleotide Archive, respectively. SRA files were converted to FASTQ files using the SRA toolkit. Samples were aligned with the kallisto pseudoalignment protocol and GRCh38 build for the human genome to produce estimated counts[10]. Treg expression values were processed using the Sleuth R Package (v0.30.0)[11].

*Cell lines and cell culture*

PY8119 cells were derived from a primary *MMTV-PyMT* tumor in the C57BL/6 background as described previously (Biswas et al., 2014). And were maintained in F12 media supplemented with 10% fetal calf serum (FCS), 10 ng $ml^{-1}$ epithelial growth factor, 2 μg $ml^{-1}$ hydrocortisone and 5 μg $ml^{-1}$ insulin. When needed, cells were detached using 2.5% trypsin at P18 and resuspended in 1:1 phosphate buffer saline: Matrigel (Corning, Corning, NY) mixture and injected in mammary fatpad at 100 μl volume per inoculation. MC38 colorectal cells were maintained in the same complete F12 media, detached using 2.5% trypsin, and resuspended in phosphate buffer saline and injected into mammary fatpad at 100 μl volume per inoculation.

*Animals*

All animals were maintained under specific pathogen-free conditions according to the IACUC guidelines. We purchased mouse sperm carrying the *Cd177* deletion allele from the UC Davis KOMP Repository and sent it to Jackson Laboratories for *in vitro* fertilization. Seven mice were obtained with 3 carrying a heterozygous deletion of *Cd177*. Mice were backcrossed to C57BL/6 for 10 generations. Genotyping of the

*Cd177*-knockout mice was performed through standard PCR procedures. More information on the generation of the *Cd177*-knockout mice is available in our previous publication[12].

*Tissue Collection and T cell suppression assay*

Fresh human breast cancer and renal cancer tumor samples were collected from patients undergoing surgical resection after informed consent, detailed above. Breast tumor samples were supplied de-identified by the Tissue Procurement Core at the University of Iowa Hospitals and Clinics and were histologically characterized by the Department of Pathology at the University of Iowa according to immunohistochemical ER, PR and HER2 biomarker staining. De-identified blood samples from healthy donors were provided by the blood bank at the University of Iowa Hospitals and Clinics. For renal cancer patients, de-identified matching blood samples were provided by the Genito-Urologic Molecular Epidemiology Resource at the Holden Comprehensive Cancer Center, University of Iowa Hospitals and Clinics. Human peripheral blood mononuclear cells (PBMCs) were isolated from whole blood using Ficoll Plaque (GE Healthcare Biosciences, Pittsburgh, PA) density gradient centrifugation or SepMate Tubes (StemCell Technologies).

Fresh tissues were directly distributed to the research laboratories after surgery, followed by enzymatic digestion and physical dissociation using gentleMACS (Miltenyi, Bergisch Gladbach, Germany) as per manufacturer's instruction. Cell suspensions were filtered through 100-μm cell strainer, magnetically enriched using anti-CD45 positive selection (Miltenyi) for tumor-infiltrating leukocytes, and immediately frozen using FBS with 5% DMSO. Breast cancer samples were used fresh or frozen without CD45 enrichment.

For *in vitro* suppression assay, naive $CD4^+$ T cells were isolated from PBMC using the human naïve $CD4^+$ T cell isolation kit II (Miltenyi, 130-094-131). TI-Tregs were labeled with anti-CD45, anti-CD4, anti-CD25, and anti-CD177 antibodies and flow-sorted and combined from 3 human breast cancer specimens to get enough number of cells, $CD45^+CD4^+CD25^+$ T cells were used as total TI-Tregs, in addition to further separation of $CD177^+$ or $CD177^-$ TI-Tregs. Naïve $CD4^+$ T cells and TI-Tregs were co-cultured at the indicated ratios (2:1, 4:1, or 8:1) on 96-well flat-bottom plates in RPMI1640 supplemented with 10% fetal bovine serum (FBS), 10 mM HEPES, $2x10^{-5}$ M 2-mercaptoethanol for 96 hr. Naïve $CD4^+$ T cells were labeled with CFSE and stimulated by anti-CD3/28 dynabeads (Thermo Fisher Scientific) and 10ng/ml IL-2 (R & D), following with flow cytometry to determine T cell proliferation indicated by CFSE dilution.

*Antibody staining and multicolor immunophenotyping of TI-Tregs*

Multicolor phenotypic panels were established using different combinations of fluorescently tagged anti-CD45 (H130), CD3 (HIT3a), CD4 (OKT4), CD25 (M-A251), CD27 (O323), CD127 (A019D5), CCR8 (433H), PD-1 (EH12.2H7), CTLA-4 (BN13), FOXP3 (206D), and CD177 (MEM-166). Cells were stained using standard immunofluorescent staining protocol and run on flow cytometry either as live cells or fixed in 4% paraformaldehyde. Antibodies were purchased from Biolegend, Molecular probes, BD Biosciences and eBioscience. Flow cytometric data was acquired on a 4-laser LSR II (BD Biosciences) and data were analyzed using FlowJo software (TreeStar, Ashland, OR). For experiments using frozen samples, cells were thawed and suspended in RPMI supplemented with 10% FBS and incubated for 1.5 hour at $37^{o}C$, 5% $CO_2$ prior to staining. Tregs were identified as $CD3^{+}CD4^{+}FOXP3^{+}$ or $CD3^{+}CD4^{+}CD25^{+}CD127^{low}$[13].

*Signature development and evaluation*

Log2 expression values from the Cancer Genome Atlas was acquired from the UCSC Xena Browser. Updated survival information was obtained from the recent work of Liu, et al[14]. After merging expression and clinical outcome data, best subset selection was performed to identify features most predictive of overall survival utilizing the leaps library in R. Feature selection was performed for three sets of genes: 1) 143 differentially-expressed genes shared between ccRCC and HCC TI-Tregs, 2) 86 genes differentially expressed in CF2 and 3) 222 genes differentially expressed in CF1.

To ensure computational feasibility of best subset selection, a custom scoring algorithm was devised. In each of the three gene sets, a random group of <30 genes was used for best subset selection, and the five genes that compromised the best five-predictor model were given a point. This process was repeated with a new random group of <30 genes until all genes had been included in exactly 500 best subset selection models. The frequency of inclusion in the top performing five-predictor model was assessed for each gene. The seven-to-eight genes that were most frequently included in the top performing five-predictor model were utilized for training subsequent machine learning algorithms. For each set of feature-selected genes, support vector machines (SVMs) were trained to predict overall survival using the e1071 (v1.7.0) R package. The 533 patients in the KIRC dataset were randomly divided into training (N=266) and testing (N=267) cohorts. SVMs were trained to discriminate overall survival status with linear kernels, and the cost-parameter was selected via cross-validation. Kaplan-Meyer curves were constructed with the survival (v2.42.6) and survminer (v0.4.3) R packages. The Cox proportion hazards regression model within the survival package was used to compute the hazard ratios between good-outcome and poor-outcome prediction groups.

*Immunohistochemistry (IHC) staining*

For IHC staining, formalin-fixed, paraffin-embedded tissue samples were sectioned at 5 μm by University of Iowa Comparative Pathology Laboratory. Sections were then de-paraffinized in a series of xylene washes and rehydrated with ethanol and water. Antigen retrieval was accomplished by immersing slides in a citrate buffer (pH 6) solution in a 110°C water bath for 15 minutes. Slides were allowed to cool-to-room temperature and were then incubated with 3% hydrogen peroxide followed by 2-5 min washes in 1X Dako buffer. Slides were then incubated with mouse monoclonal anti-CD177 antibody (clone C4C; Sigma) for 1 hours and then Dako Mouse Envision System HRP for 30 minutes. The slides were developed with 3,3'-diaminobenzidine (Dako DAB plus) for 5 min followed by 3-minute incubation in 3,3′-diaminobenzidine tetrahydrochloride (Dako DAB enhancer). Samples were then counterstained with hematoxylin, rinsed in water, cleared in xylene and then mounted with Permount. Histological evaluations were performed by comparative pathologist Dr. Katherine Gibson-Corley.

*Statistical Analysis*

Statistical Analyses were performed in R (v3.5.1). Two-sample significance testing utilized Welch's T test, with significance testing for more than three samples utilizing one-way analysis of variance (ANOVA) with Tukey honest significance determination for correcting multiple comparisons. Differential gene expression for single-cell RNA-seq data was performed in the Seurat R package using the Wilcoxon rank sum test with p-values adjusted using the Benjamini-Hochberg method[2,3]. Differential gene expression of bulk RNA-sequencing Treg expression data utilized Wald testing in the Sleuth R Package[11]. Differential gene expression between branches of the cell trajectory manifold was performed in the Monocle R package using likelihood ratio testing[4].

*Data Availability*

Quantified gene expression counts and V(D)J T cell receptor sequences for SCRS are available at the Gene Expression Omnibus (GEO) at GSE121638.

### Supplementary Figure Legends

**Supplementary Figure 1. Differential gene expression of TI-Tregs in single-cell RNA-seq and pooled RNA-seq.**

(a). tSNE projection of T cells from five HCC patients with normal PB cells and TI cells. TI Treg population (green, n=634) and PB Treg population (grey, n=264) were isolated for further analysis. (b). tSNE projection with highlighted expression of Treg markers, FOXP3 and IL2RA (CD25). (c). Differential gene expression analysis using the log2-fold change expression versus the difference in the percent of cell expressing the gene comparing TI versus PB Tregs (Δ Percentage Difference). (d). Venn diagram of the overlap in differentially-expressed tumor-infiltrating Tregs compared to peripheral-blood Tregs in breast carcinoma (GSE89225), colorectal adenocarcinoma (PRJEB11844), and non-small cell lung squamous carcinoma (PRJEB11844). (e). Heatmap differential genes shared between the three datasets with immune-related genes labeled. Genes are displayed in Log2 fold-change (FC) comparing tumor-infiltrating Tregs versus peripheral-blood Tregs with FDR q-values <0.05.

**Supplementary Figure 2. Transcriptional Heterogeneity in single-cell sequencing of HCC Tregs.**

(a). Trajectory manifold of Tregs from the HCC using the Monocle 2 algorithm. (b). Pseudo-time projections of transcriptional changes in immune genes based on the manifold. (c). Cell trajectory projections of transcriptional changes in immune genes based on the manifold. $\bar{x}$ denotes the scaled mean of each pole of the manifold. (d). Gene signature analysis of the poles of the trajectory manifold. *** $P < 0.001$, **** $P < 0.0001$. (e). Results of the cell cycle regression analysis of single cells for each cell fate using the Seurat R package.

**Supplementary Figure 3. Clonotype analysis of ccRCC-infiltrating and peripheral-blood Tregs.**

(a). Percentages of assigned clonotypes by patients in peripheral-blood (PB, upper panel) and tumor-infiltrating Tregs (TM, lower panel). (b). Relative increase in clonotypes with two copies in the same patient comparing peripheral-blood (grey) to tumor-infiltrating (orange). Significance testing utilized T test with Welch's correction. (c). Relative increase in clonotypes with three or more copies in the same patient comparing peripheral-blood (grey) to tumor-infiltrating (orange) Tregs. Significance testing utilized T test with Welch's correction. (d). Cell trajectory projections of shared clonotype percentages by cell fate, combining the peripheral-blood and tumor-infiltrating clonotypes of a single patient. Significance based on $\chi^2$ testing.

**Supplementary Figure 4. $CD177^+$ tumor-infiltrating lymphocytes are comprised of Tregs.** (a-b). Representative flow cytometry data gating on Tregs identified as $CD25^+CD127^-$ (a) or $FoxP3^+$ (b) and isolated from TI in breast cancer tissue or PBMCs. (c). IHC staining protocol is established and the specificity of the anti-CD177 antibody is confirmed using isotype IgG on sequential sections of normal human colon tissue, scale bar = 100μm. (d). CD177 IHC staining on different normal human tissues showed CD177 expression on normal epithelium for select tissues as well as infiltrating leukocytes. Black arrows: epithelial cells; orange arrows: lymphocytes scale; bar = 200μm. (e). Representative dual IHC staining for CD177 (red) and FoxP3 (brown) in breast carcinoma section. Dual-positive cells are indicated with red arrows and FoxP3-positive cells are indicated by black arrows.

**Supplementary Figure 5. Suppressive phenotype of CD177$^+$ Treg in human and mouse assays.** (a). CD177$^+$ TI-Tregs are suppressive and inhibit CD8 T cell proliferation. Total, CD177$^+$, or CD177$^-$ TI-Tregs were purified from fresh human breast cancer specimens (combined from 3 patients) using flow cytometry and co-cultured with naïve CD8$^+$ T cells from PBMC for *ex vivo* suppression assay at the indicated ratios. (b). Tregs isolated form CD177 WT tumor bearing mice are significantly more suppressive than KO Tregs (unpaired Welch's t-test), n=3

**Supplementary Figure 6. Survival predictions based on Treg single-cell gene signatures across the TCGA cohort.** (a). Disease-specific survival prediction with Cox proportional hazard ratio and -log10(P-value) based on log-rank testing across the 24 largest TCGA datasets using the TI-Treg signature. (b). Disease-specific Survival prediction with Cox proportional hazard ratio and -log10(P-value) based on log-rank testing across the 24 largest TCGA datasets using the Cell Fate #1 Treg signature. (c). Disease-specific survival prediction with Cox proportional hazard ratio and -log10(P-value) based on log-rank testing across the 24 largest TCGA datasets using the Cell Fate #2 signature.
