## Supplementary material for "Transcriptional heterogeneity in cancer-associated regulatory T cells is predictive of survival"

Supplemental Figure 1

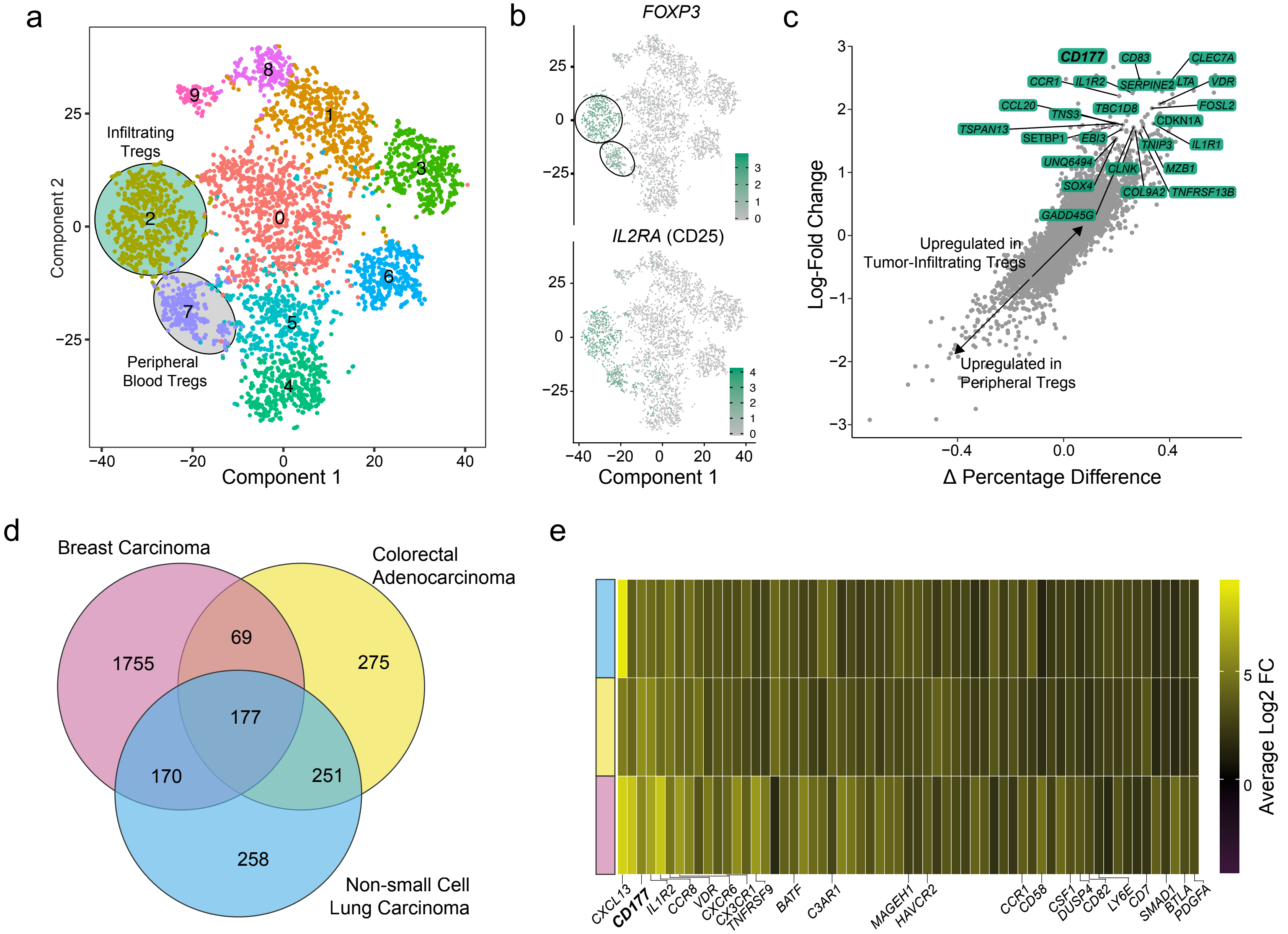

### Supplemental Figure 2

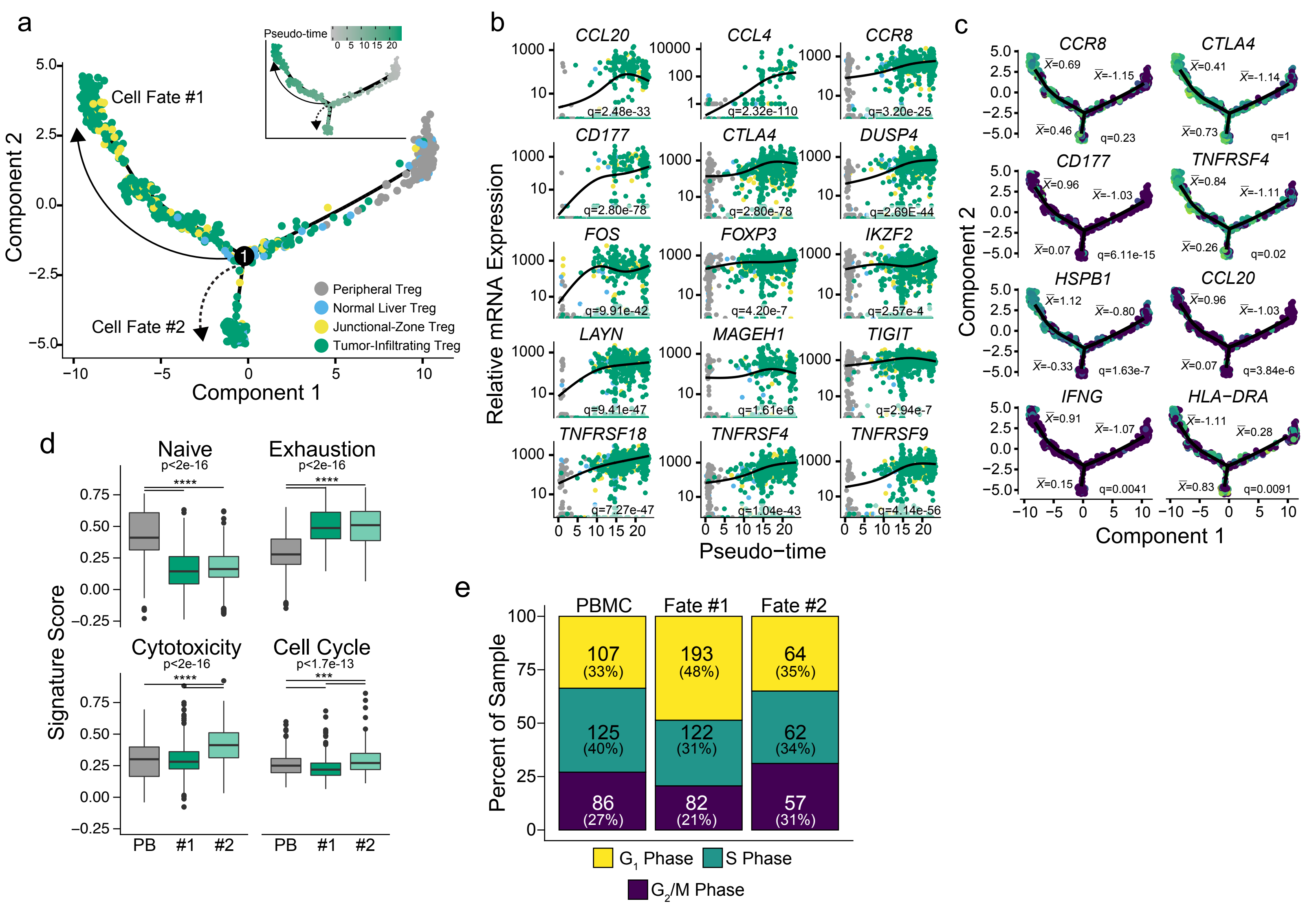

### Supplemental Figure 3

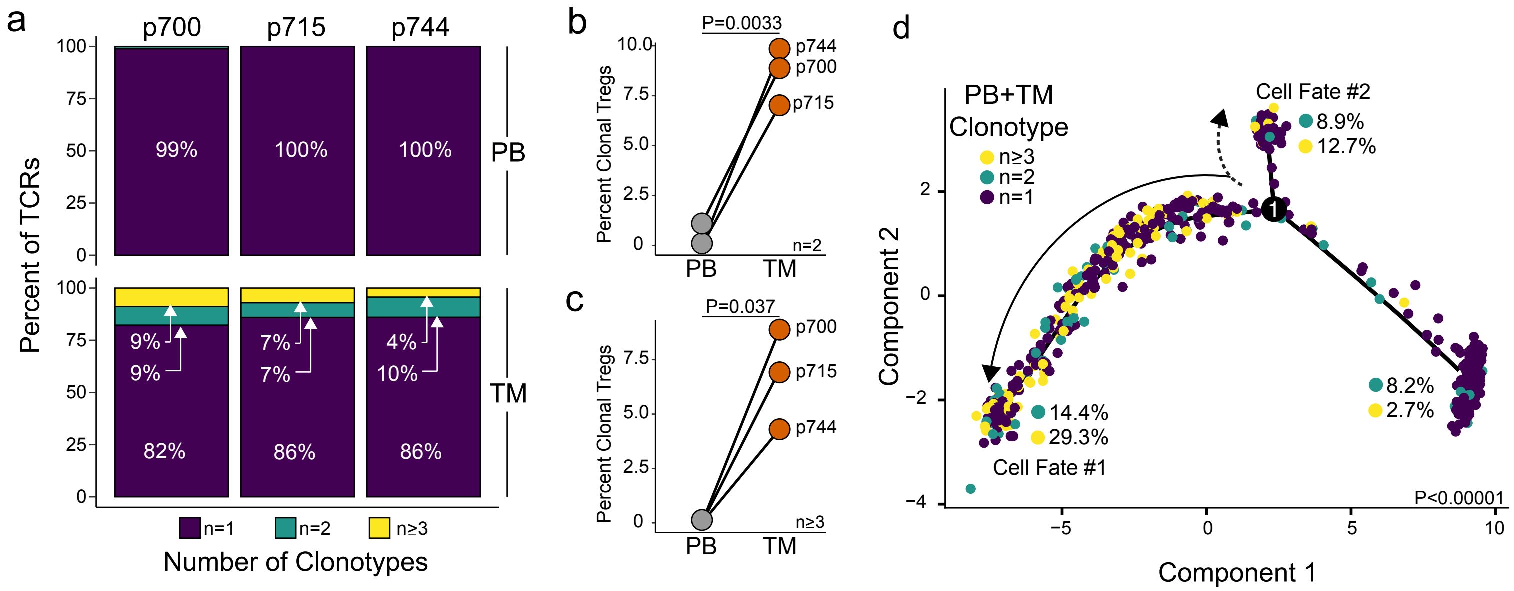

Supplemental Figure 4

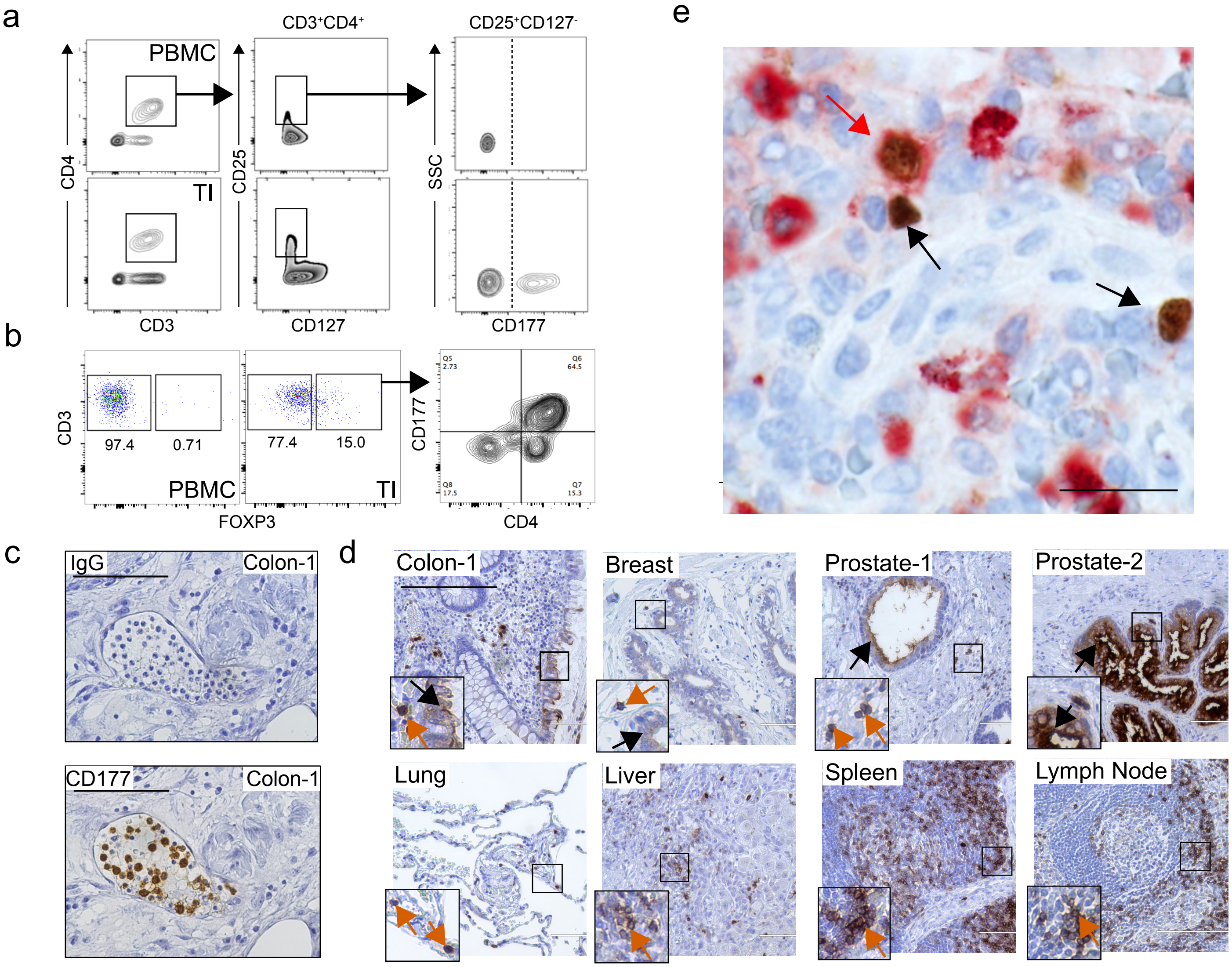

Supplemental Figure 5

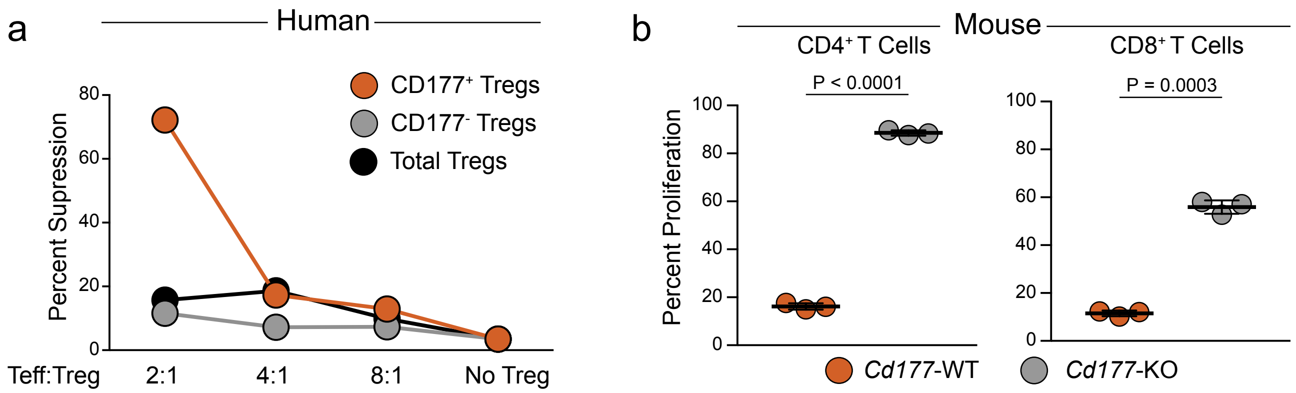

### Supplemental Figure 6

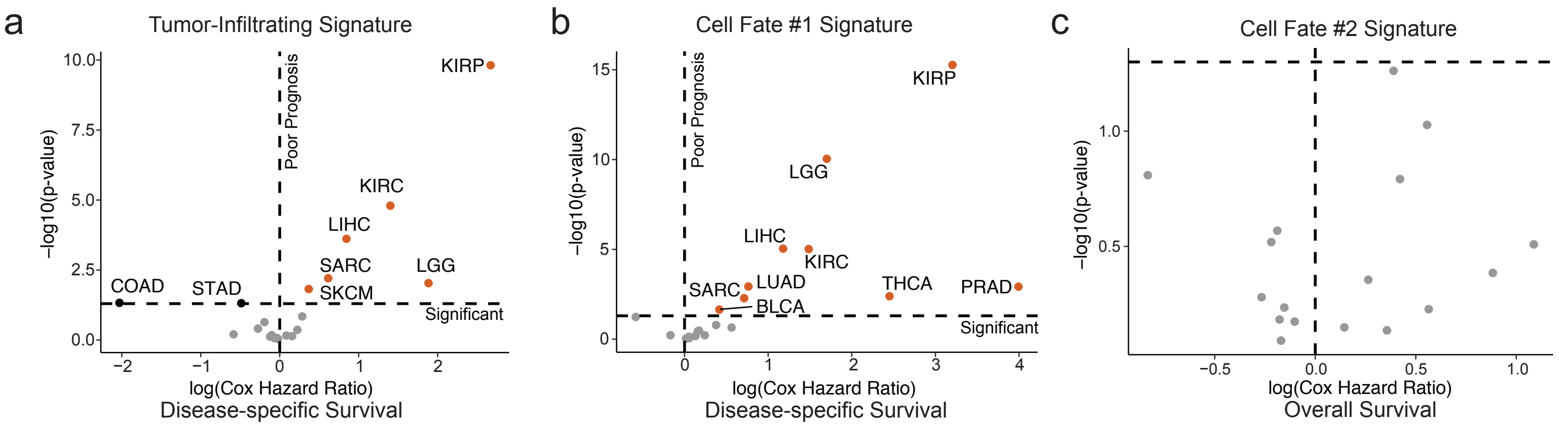
